# Recovery of a minor cryo-EM particle population reveals conformational equilibria linking cofactor loading, turnover, and reactivation in methionine synthase

**DOI:** 10.64898/2026.07.27.741092

**Authors:** Haoyue Wang, Maxwell B. Watkins, Pasa Suksmith, Nozomi Ando

**Author notes:** Correspondence should be addressed to **Corresponding Author: Nozomi Ando,**. These authors contributed equally (Maxwell B. Watkins, Pasa Suksmith).

## Abstract

Cobalamin-dependent methionine synthase (MetH) is a highly dynamic enzyme that requires large-scale domain rearrangements to facilitate turnover and reactivation. Despite significant biochemical and structural efforts, a full picture of the conformational changes underlying the transition between MetH turnover and reactivation has remained elusive. Here, we leverage state-of-the-art cryo-electron microscopy (cryo-EM) methods to reanalyze an old dataset and gain new insights into these changes. By utilizing a neural-network particle picker, performing careful classification of particles, and incorporating high-resolution information to determine initial particle orientations, we uncovered and reconstructed a minor population of reactivation-state MetH (∼4.5-5 Å resolution) from a dataset that is dominated by resting-state MetH. This structure represents the first fully intact structure of the enzyme with a physiological cobalamin cofactor bound. These findings highlight the potential for improved cryo-EM methodology to uncover previously hidden details about enzyme function and deepen our understanding of structural ensembles.

---

Cobalamin-dependent methionine synthase (MetH) catalyzes methyl transfer from 5-methyltetrahydrofolate (CH_3_-H_4_folate) to L-homocysteine (Hcy) to produce L-methionine (Fig. 1). During turnover, the vitamin B_12_ cofactor (cobalamin) acts as an intermediate methyl carrier, alternating between the CH_3_-Cob(III) (methylcobalamin) and Cob(I) oxidation states (Fig. 1B). Cob(I) is highly oxygen-sensitive and occasionally oxidized to an inactive Cob(II) state. The cofactor is reactivated by reductive methylation using an electron donor (e.g., flavodoxin in *E. coli*^1^) and *S*-adenosylmethionine (AdoMet) as the methyl donor (Fig. 1B). The three substrates and the cofactor bind to four separate domains of MetH: the Hcy, Folate, B_12_, and AdoMet domains, from N- to C-terminus (Fig. 1A). Throughout turnover and reactivation, the cofactor must interact with three spatially separated active sites, necessitating significant conformational rearrangements.

**Figure 1.**
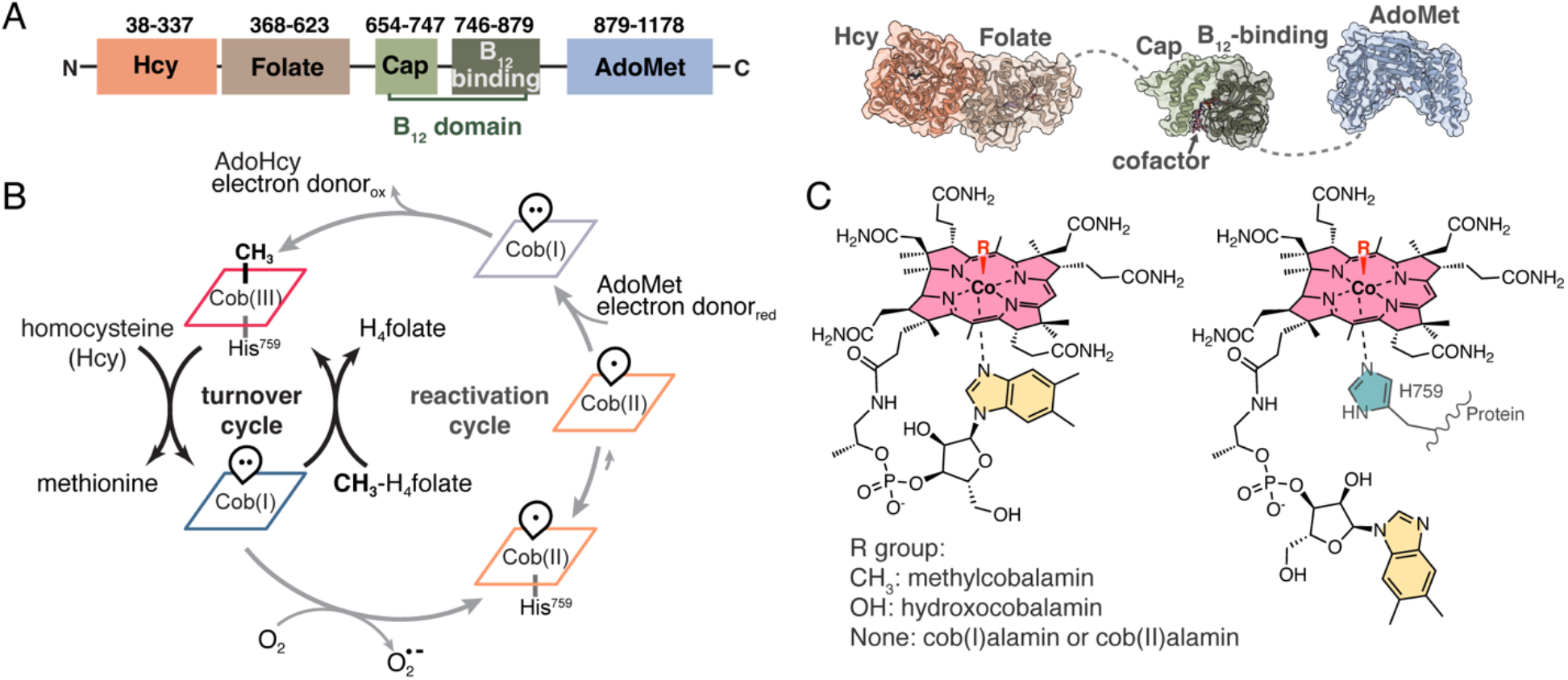
Overview of MetH. (A) From N- to C-terminus, the four domains of MetH bind homocysteine (Hcy), 5-methyltetrahydrofolate (CH3-H4folate), cobalamin (B12), and *S*-adenosylmethionine (AdoMet). The flexibly linked B12 domain contains a mobile protective cap subdomain and the cofactor-binding subdomain. Domain boundaries are shown in *T. filiformis* numbering from Uniprot. (B) During turnover (black arrows), MetH cycles between the CH3-Cob(III) and Cob(I) states to produce to methionine and H4folate from Hcy and CH3-Cob(III). Oxidation of Cob(I) to an inactive Cob(II) state requires reactivation by AdoMet and an electron donor (gray arrows). (C) Cobalamin can coordinate different upper and lower ligands. In free cobalaminm the lower ligand is a dimethylbenzimidazole (DMB) tail, but in MetH, the lower ligand is replaced by a conserved histidine (His759 in *E. coli* and *T. filiformis* numbering). The “His-on” state is defined by His759 coordinating the central cobalt ion.

The dynamic nature of MetH has made it challenging to characterize structurally. Early limited proteolysis experiments showed that MetH follows distinct degradation pathways during turnover and reactivation and that, in the Cob(II) state, the pathway further depends on whether a conserved histidine (His759 in *Escherichia coli* and *Thermus filiformis* numbering) serves as the lower ligand of cobalamin (“His-on”, Fig. 1C) or dissociates from it (“His-off”).^2^ Additional studies suggested that MetH adopts at least four conformations.^3^ However, for decades, only structures of MetH fragments were available.^4–11^ The first multidomain structures of the C-terminal half fragment of *E. coli* MetH revealed the so-called reactivation conformation, where the B_12_ domain interacts with the active site of the AdoMet domain in a “cap-off” state, exposing the upper face of the cobalamin cofactor for methyl transfer.^6,9,10^ By contrast, for many years, the only available structure representing the turnover cycle was the isolated B_12_ domain bound to methylcobalamin.^4^ This structure captured a “cap-on” conformation (PDB: 1BMT), with the cofactor sandwiched between the two B_12_ subdomains, shielding it from unwanted reactivity.

Recently, new structural data on full-length MetH have provided a more holistic understanding of the enzyme’s conformational landscape. In 2023, we used solution small-angle X-ray scattering (SAXS) to show that the physiologically relevant oxidation states share a similar conformational ensemble in the absence of substrates.^12^ Cryo-EM of *T. filiformis* MetH, together with SAXS data, further suggested that this shared ensemble primarily contains a stable “resting state” conformation in which a cap-on B_12_ domain is sequestered between the Hcy and Folate domains, while the flexibly linked AdoMet domain remains mobile and unresolved (PDB: 8G3H, EMD-29699).^12^ That same year, a full-length crystal structure of apo-MetH from *Thermus thermophilus* was reported (PDB: 8SSC).^13^ The C-terminal half of this structure adopts the reactivation conformation previously observed in fragment structures^6,9,10^ and packs against the N-terminal half through a ∼900 Å^2^ interface between the AdoMet and Hcy domains – an interaction that was interpreted to be transient. In 2024, additional important structures were reported using constructs of *T. thermophilus* MetH lacking the AdoMet domain, including one that recapitulated the cap-on resting-state conformation seen in 8G3H and other conformations relevant for the turnover cycle.^14^ In 2026, full-length cryo-EM structures of human methionine synthase (MTR) captured the C-terminal half in a reactivation conformation in both the cofactor-less apo state and bound to a non-physiological hydroxocoblamin ligand. In the latter, the enzyme exists in a His-on (PDB: 9SSS) and His-off (PDB: 9SST) equilibrium,^15^ consistent with spectroscopic data from *E. coli* MetH.^16^

Despite these advances, the structural dynamics linking MetH turnover and reactivation remain unclear. Based on our structural results^12^ and published EPR data,^16^ we previously proposed that Cob(II) MetH in the absence of substrates interconverts largely between the dominant cap-on resting-state conformation and a minor cap-off reactivation-state conformation, but we did not directly observe the latter in our cryo-EM dataset. Here, we leveraged improvements in cryo-EM data processing and methodology to reprocess the original dataset that gave rise to the resting-state structure of *T. filiformis* MetH and identified a rare population of only 22.5k particles, from which we reconstructed a stable full-length reactivation conformation at an estimated resolution of ∼4.5-5 Å. This minor conformation recapitulates the *T. thermophilus* apo-MetH structure (PDB: 8SSC) that was previously proposed to be transient and matches our solution SAXS data on the apo *T. filiformis* enzyme (described below), revealing an unexpected connection between the inactive Cob(II) state of MetH and the cofactor-less apo state.

We began by reprocessing the original raw data that yielded our previous cryo-EM resting-state map of *T. filiformis* Cob(II) MetH (EMD-29699) using a new workflow in CryoSPARC^17^ (Fig. 2 and Supplementary Information). Improved particle picking, contrast transfer function (CTF) refinement with beam-shift correction,^18^ and particle polishing^19^ slightly improved the quality of the map, yielding a smoother Fourier shell correlation (FSC) curve and a 0.2 Å improvement in overall nominal resolution (Fig 2A,C), although map anisotropy was still present. High-resolution heterogeneous *ab initio* reconstruction (HR-HAIR)^20^ with a maximum resolution cut-off of 3.2 Å further removed particles that align to high resolution in 2D classes but ultimately degrade the reconstruction. Subsequent local refinement yielded a much higher-quality final map with significantly more isotropic density than our original map (Figs. 2C, S1A, S2). Non-uniform (NU) refinement^21^ was also tested but produced a map with greater anisotropy with no gain in resolution (Fig. S1B), as also observed in the original HR-HAIR paper.^20^

**Figure 2.**
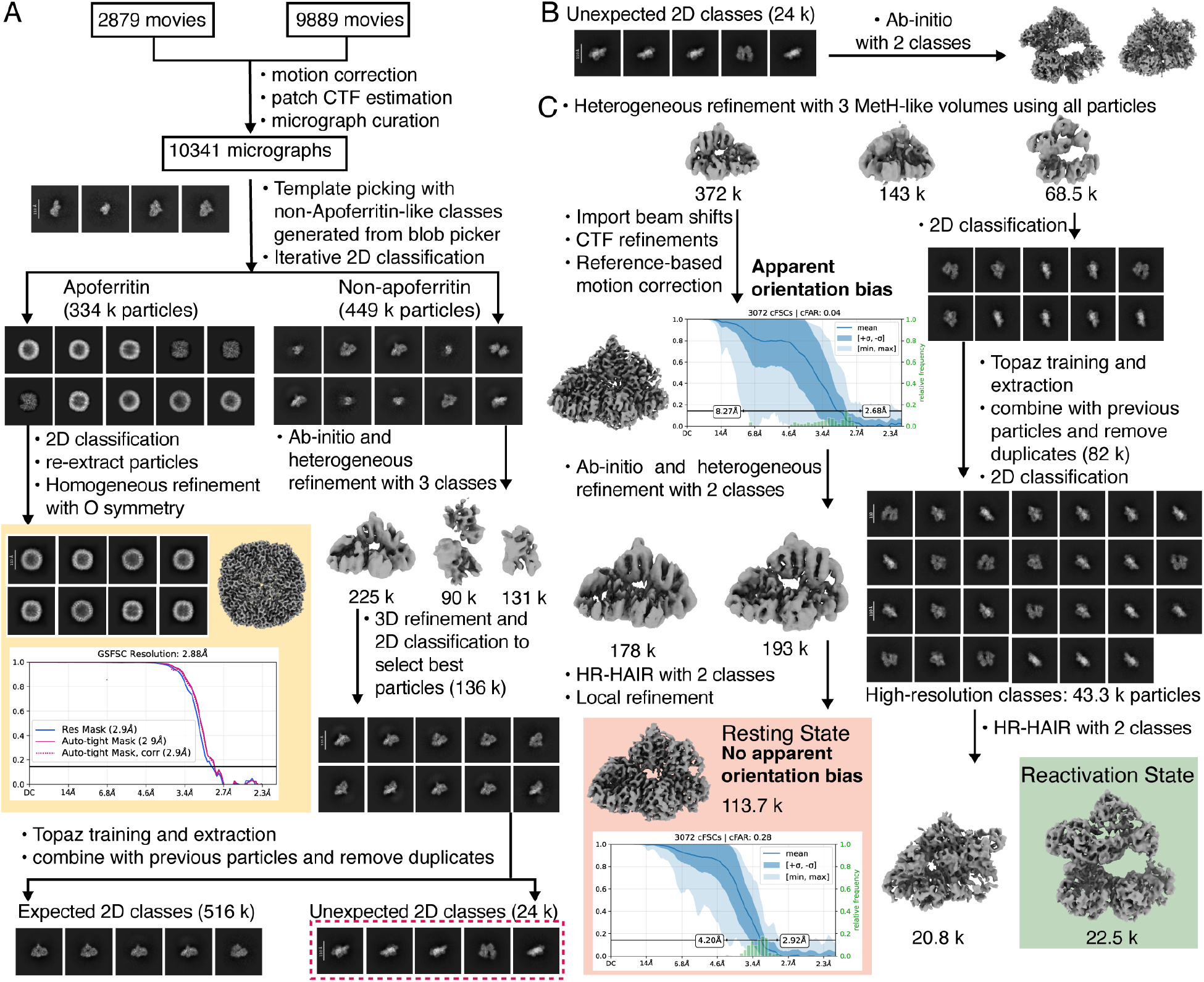
The conformational ensemble of *T. filiformis* Cob(II) MetH contains a predominant resting state and a minor reactivation state. Reprocessing of a previously published cryo-EM dataset12 co-frozen with 2 μM apoferritin in 50 mM HEPES, 150 mM NaCl, 2.5 mM DTT (pH 7.6). (A) Conventional cryo-EM processing yielded only the dominant resting-state conformation, while Topaz-based particle picking and extensive 2D classification uncovered a rare conformational population. (B) Initial *ab initio* reconstruction yielded starting volumes, including one consistent with the reactivation conformation. (C) High-resolution heterogeneous *ab initio* reconstruction (HR-HAIR) further filtered junk particles, yielding a higher-quality resting-state map (red box) and a map of the rare full-length reactivation conformation (green box).

Under the conditions used to obtain the original cryo-EM dataset (50 mM 4-(2-hydroxyethyl)-1-piperazineethanesulfonic acid (HEPES), 150 mM NaCl, 2.5 mM dithiothreitol (DTT), pH 7.6), the cofactor is expected to be predominantly 5-coordinate His-on Cob(II), with a UV-visible absorbance peak at 477 nm.^12^ Consistent with this expectation, the corrin ring in our new resting-state reconstruction aligns well with that observed in previous “cap-on, His-on” structures^4,14^ rather than in “cap-off, His-off” structures where the corrin ring has swung away from the B_12_ domain.^6,9,10^ However, the reconstruction does not show clear evidence for a fully typical His-on geometry: we do not observe density connecting His759 to the cobalt ion, and the refined His-Co bond distance (3.0 Å) is longer than in previously reported His-on MetH structures^4,14^ (typically <2.5 Å) (Fig. S3). These observations suggest that elongation or weakening of the His-Co interaction does not necessarily require a large displacement of the corrin ring.

To our surprise, Topaz-based particle picking^22^ trained on high-resolution particles from the resting-state reconstruction, followed by thorough 2D classification (200 classes, 80 iterations, batch size of 400 per class for 1.8M picked particles), revealed previously unseen 2D classes (Fig. 2A, red box). *Ab initio* reconstruction and heterogeneous refinement showed that these particles reconstruct a conformation resembling the reactivation state (Fig. 2B). Topaz was then retrained using high-resolution particles from this initial reconstruction to enrich this rare population. HR-HAIR with two classes and a 3.4 Å maximum resolution cut-off yielded a junk class and a reconstruction of a full-length, cap-off reactivation conformation (Figs. 2C, S4A). The 2D classes of the particles retained after HR-HAIR show clearly recognizable front views of the reactivation conformation (Fig. S5A), while many of the junk classes contain ambiguous views (Fig. S5B).

Following HR-HAIR, we also performed local and NU refinements using a 12 Å initial low-pass filter, which yielded maps with gold-standard FSC resolutions of 4.6 and 5.1 Å, respectively (Fig. S4B-C). Although the FSC calculated from half-maps reconstructed using HR-HAIR-derived particle poses yielded a nominal resolution of 3.8 Å, sharpening did not reveal corresponding high-resolution features above noise. Visually, the HR-HAIR map is closer to ∼4.5-5 Å resolution, consistent with the estimates from NU and local refinement. This discrepancy is attributed to the lack of independent half-set refinement during HR-HAIR. Nonetheless, all three maps were highly similar overall (Fig. S4) and closely matched the AlphaFold3 prediction of the full-length *T. filiformis* MetH. Because the unsharpened HR-HAIR map showed better-defined density, it was used for restrained model refinement in ISOLDE (Supplemental Information). This final model is highly similar to a more conservative model obtained by per-domain rigid-body docking of the AlphaFold3 prediction into the local-refinement map (C? RMSD of 0.596 Å of all pairs), but incorporates ligands and shows improved Q-scores (Fig. S6).

The final reactivation-state map was reconstructed from only 22.5k particles, compared with 372k before HR-HAIR and 113.7k after further curation for the final resting-state map (Fig. 2C). Nonetheless, this represents the first full-length reactivation-state structure with a physiological cofactor (in this case, Cob(II)), and the resolution is sufficiently high to build a quasi-atomistic model and draw several important conclusions. The N-terminal half adopts a rigid conformation conserved in all MetH structures, while the C-terminal half closely resembles the reactivation conformation seen in fragment structures of *E. coli* MetH (Ca RMSD of 1.28 Å over 225 pruned atom pairs relative to PDB: 3BUL). Strong corrin-ring density is observed, allowing the cofactor to be docked by aligning the B_12_-binding subdomain with previous structures. Interestingly, as in our new resting-state map, the density of the corrin ring aligns better with the position observed in the “cap-on, His-on” structure 1BMT than with that in the “cap-off, His-off” structure 3BUL, where the corrin ring swings out (Fig. 3A). However, the resolution of this map is insufficient to resolve the His759 side chain, precluding unambiguous assignment of the His-on/off state.

**Figure 3.**
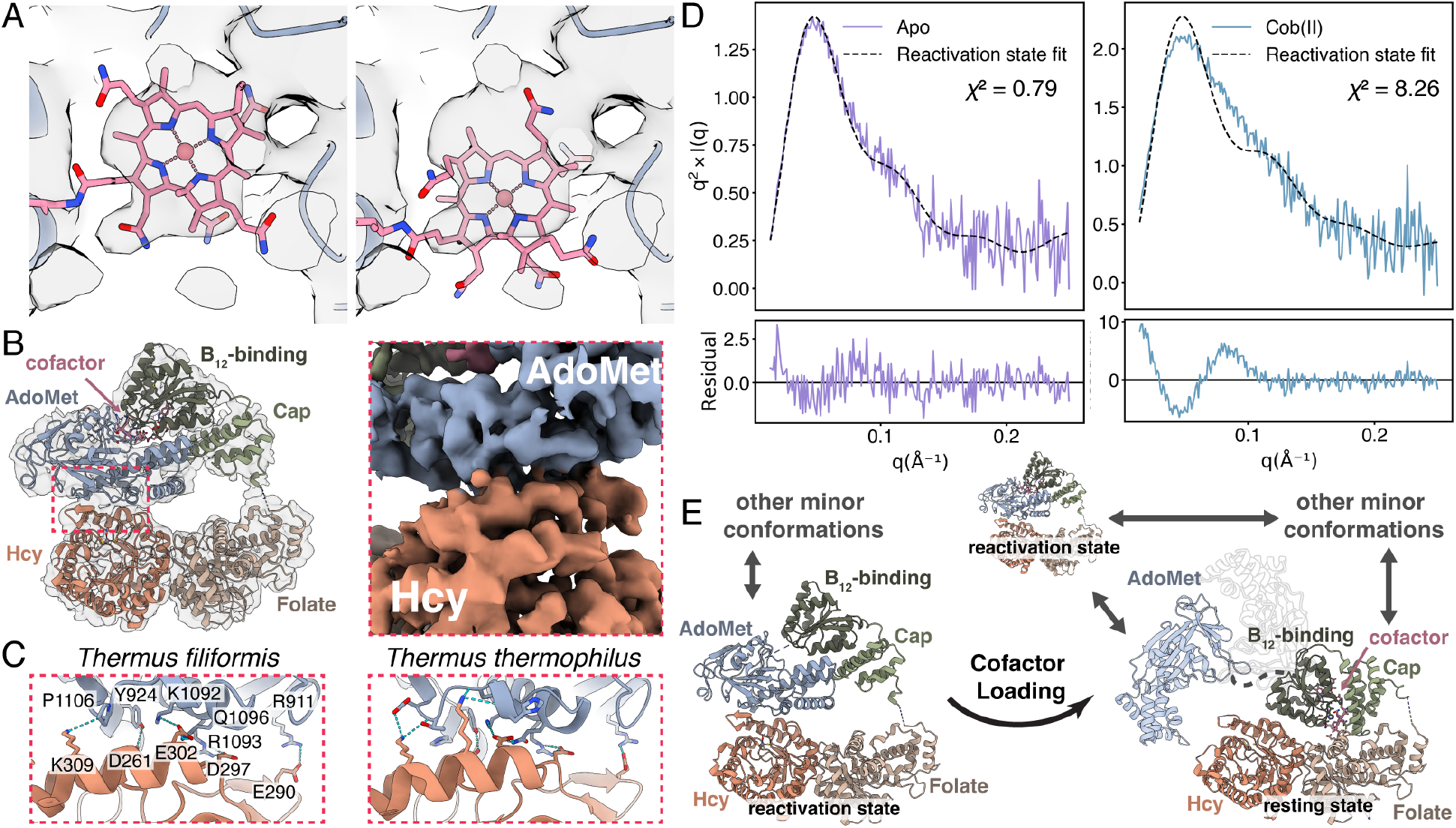
The reactivation state links cofactor loading to the MetH conformational cycle.. (A) Rigid-body alignment of the B12-binding domain with previous His-on (PDB: 1BMT, left) and His-off (PDB: 3BUL, right) structures shows that the corrin-ring position of the former better fits the density. (B) Cryo-EM model of the full-length cap-off reactivation conformation contains a stable AdoMet-Hcy domain interface. (C) This interface resembles that in the *T. thermophilus* apo-MetH crytal structure (PDB: 8SSC). (D) The reactivation-state cryo-EM model agrees with SAXS data collected on *T. filiformis* MetH in the apo state (left) but not the Cob(II) state (right). SAXS was performed in the same buffer as cryo-EM (50 mM HEPES, 150 mM NaCl, 1-2.5 mM DTT, pH 7.6). (E) Proposed model for the conformational distribution of *T. filiformis* MetH. In the absence of cofactor, MetH is predominantly in the cap-off reactivation state. Cofactor loading shifts the distrbution to a cap-on resting state that interconverts with a minor reactivation-state population, particularly in cofactor oxidation states where the Co-His interaction is weakened. In the resting state, the three N-terminal domains form a stable core whereas the AdoMet domain (light blue) remains mobile.

Another unexpected finding was a stable interface between the Hcy and AdoMet domains (Fig. 3B), similar to that previously observed in *T. thermophilus* apo-MetH (PDB: 8SSC, Fig. 3C). *QtPisa*^23–25^ estimated both interfaces as moderately strong (∼-10 kcal/mol), supported by a large number of non-electrostatic interactions and several electrostatic interactions (Fig. S7A). The strength of the interface may reflect the delicate balance needed between stabilizing the reactivation state and enabling structural transitions to other states. The residues at the interface are relatively conserved between *T. filiformis* and *T. thermophilus* (Fig. 3C). Across a broader set of 2,205 representative MetH sequences, moderate conservation is observed for residues involved in non-electrostatic interactions (Fig. S7B, gray), while those involved in electrostatic interactions are less conserved (Fig. S7B, colored). The Hcy-AdoMet interface in the reactivation state partially overlaps with the Hcy-B_12_-subdomain interface in the resting state, indicating that interconversion requires disengagement of the B_12_-binding subdomain from the Hcy domain to accommodate the AdoMet domain, together with substantial rearrangement and uncapping of the B_12_ domain (Fig. S8).

Interestingly, the full-length reactivation conformation resolved by cryo-EM fits the solution SAXS profile of *T. filiformis* apo-MetH reasonably well but clearly does not fit that of Cob(II) *T. filiformis* MetH (Fig. 3D), consistent with it being a minor species in the Cob(II) cryo-EM dataset. Together with previous structures of apo-MetH,^13^ our results suggest that before cofactor loading, the enzyme is predominantly in a stable cap-off reactivation state, whereas cofactor loading shifts the conformational ensemble towards a predominantly cap-on resting state in equilibrium with a small but detectable population of the reactivation state (Fig. 3E).

The presence of a small reactivation-state population in Cob(II) MetH is consistent with biochemical observations that only a small fraction of the Cob(II) enzyme is poised for reactivation in *E. coli*.^2^ Limited proteolysis and spectroscopic data suggested that ∼15% of as-isolated *E. coli* MetH is His-off^16^ and that the His-on-to-off transition favors the reactivation conformation.^2^ Flavodoxin binding likewise favors the His-off state,^1^ and we previously showed that flavodoxin selectively binds Cob(II) MetH despite its predominantly resting-state conformational ensemble, similar to other physiologically relevant oxidation states.^12^ Based on these observations, we had proposed that the Cob(II) ensemble contains a minor His-off reactivation conformation recognized by flavodoxin, thereby siphoning the enzyme from the turnover cycle into the reactivation cycle.^12^ While the resolution of our Cob(II) reactivation-state map is insufficient to unambiguously assign the His-on/off state, our refined resting-state model displays an elongated His-Co distance despite the corrin ring occupying a position similar to that observed in previous His-on structures with much shorter His-Co distances.^4,14^ Therefore, a similar corrin-ring position in the reactivation state may still be consistent with a weakened His-Co interaction. A model in which a weakened His-Co interaction promotes sampling of the reactivation conformation would also help explain why both the apo enzyme,^13^ which lacks a cobalamin cofactor, and the non-physiological hydroxocobalamin-bound enzyme,^15^ which has previously been observed in a His-on/off equilibrium, adopt this conformation. Structural studies of human MTR have suggested that the apo enzyme enters the reactivation cycle upon cofactor loading, a process that involves eukaryote-specific accessory proteins.^15^ While the *in vivo* mechanism of cofactor loading remains unclear in bacteria, it is known that certain bacterial MetHs, including the *T. filiformis* and *T. thermophilus* enzymes, can be reconstituted *in vitro* from the apo form,^12,13^ and our SAXS data suggest that bacterial MetHs also adopt a stable reactivation conformation to facilitate cofactor loading (Fig. 3D). Together, these observations suggest that transition to the reactivation conformation may represent a common mechanism by which MetH responds to the absence or loss of a catalytically competent cobalamin state. This conformation may represent a state in which movement of the cobalamin cofactor, as required for both cofactor loading and methyl transfer from AdoMet, is less restricted.

Through advances in cryo-EM data processing, we resolved a minor population of reactivation-state particles that had been buried within a dataset dominated by resting-state particles. Its recovery provides direct structural evidence that these conformations interconvert within the Cob(II) MetH ensemble. This conformational equilibrium is modulated by the presence and chemical state of the cobalamin cofactor, linking cofactor loading, turnover, and reactivation in MetH. As experimental and computational methods continue to improve, additional low-population conformations will likely emerge, providing an increasingly complete picture of the structural ensemble that underlies MetH catalysis and reactivation.

## Supporting information

per-domain-docking-model

## Supporting Information

Additional methods, figures, and tables are available in the SI. The rigid-body docking model of the reactivation state is provided as a supplementary coordinate file.

## Author Contributions

H.W. performed the majority of data analysis; M.B.W. collected the original cryo-EM data and built the new resting-state model; P.S. built the reactivation-state model; N.A. supervised the project and acquired funding. All authors contributed to writing and approved the final manuscript.

## Data Availability

Cryo-EM maps and models are available as EMD-78313/PDB 37MP (resting state) and EMD-78314/PDB 37MQ (reactivation state). The locally refined reactivation-state map is available as EMD-79288. Raw EM data are available as EMPIAR-13818.

## Acknowledgments

We thank Jon Cohen for assistance with manuscript editing. This work used a Xenocs BioXolver funded by NIH S10OD028617 and the National Center for CryoEM Access and Training (NCCAT) and the Simons Electron Microscopy Center located at the New York Structural Biology Center, supported by the NIH Common Fund Transformative High Resolution Cryo-Electron Microscopy program (U24 GM129539, and NIGMS R24 GM154192) and by grants from the Simons Foundation (SF349247) and NY State Assembly. P.S. was supported by NIH training grant T32GM138826. This work was supported by an NIH grant GM124847 to N.A.

