## Supplementary Information for "Recovery of a minor cryo-EM particle population reveals conformational equilibria linking cofactor loading, turnover, and reactivation in methionine synthase"

‡ These authors contributed equally

**This PDF file includes:**

Methods

Tables S1-S2

Figures S1-S8

Supporting References

### Methods

#### Cryo-EM Data Processing

Raw movies that yielded our previously published *Thermus filiformis* Cob(II) MetH reconstruction (EMD-29699; two datasets collected on identically prepared grids)[1] were reprocessed. For the original data collection, *T. filiformis* Cob(II) MetH was mixed with horse spleen apoferritin, both in 50 mM 4-(2-hydroxyethyl)-1-piperazineethanesulfonic acid (HEPES), 150 mM NaCl, 2.5 mM dithiothreitol (DTT), pH 7.6, at final concentrations of 2 μM each. Raw movies were combined and imported into CryoSPARC 5[2] for patch motion correction and patch contrast transfer function (CTF) estimation. Micrographs were curated to a subset of 10,341 of the original micrographs. Non-apoferritin-like templates were generated by 2D classification from particles picked using blob picker with a radius between 80 and 100 Å. Template-picked particles were extracted with a 256-pixel box size and Fourier-cropped by a factor of 2. Iterative 2D classification was performed to select non-apoferritin particles, prior to *ab initio* reconstruction and heterogeneous refinement with 3 classes. Particles from the class resembling MetH were re-extracted without Fourier cropping for *ab initio* reconstruction and heterogeneous refinement with 2 classes, followed by a round of 2D classification to select the best particles. These particles were used to train a Topaz model for repicking particles.[3]

Topaz training was performed using the default parameters. Topaz-picked particles were extracted and Fourier-cropped by a factor 2 to perform one round of 2D classification. High-resolution classes were reextracted without cropping and were combined with the template-picked particles, with duplicate particles removed (40 Å minimum separation distance), to yield 564,934 particles. 2D classification with 100 classes, 40 iterations, and batch size of 400 per class was performed, resulting in new unexpected 2D classes. *Ab initio* reconstruction with two classes was performed on the unexpected 2D classes to generate reference volumes. All non-apoferritin particles aligning to high-resolution were selected for heterogeneous refinement using the resting-state map and the two newly generated *ab initio* volumes. Particles assigned to resting-state or reactivation-state maps were separated for downstream processing.

For the resting-state particles, beam-shift parameters were imported after the initial non-uniform refinement[4] and incorporated into subsequent global and local CTF refinement[5] using 25 optics groups. One round of reference-based motion correction[6] followed by additional global and local CTF refinement was performed before a final non-uniform refinement, yielding a 3.3 Å resolution map from 371,160 particles with apparent orientation bias. The particles were then filtered by one round of *ab initio* and heterogeneous refinement, and the more isotropic class was further classified by HR-HAIR with 2 classes and a maximum resolution cut-off of 3.2 Å. The final map, with a resolution of 3.4 Å estimated by gold-standard Fourier shell correlation (FSC), was obtained by local refinement of the more isotropic class (Fig. S1A). Non-uniform (NU) refinement of the same particles yielded a map of comparable nominal resolution but greater anisotropy (Fig. S1B).

Because of the limited number of reactivation-state particles, additional 2D classification was performed to generate templates for a second round of Topaz picking and particle extraction. Newly picked particles were merged with the previous particle stack, and duplicates (100 Å minimum separation distance) were removed. A subsequent round of 2D classification identified 43,324 particles belonging to high-resolution 2D classes. High-resolution heterogeneous *ab initio* reconstruction (HR-HAIR)[7] was then performed using 2 classes. Maximum resolution cutoffs of 3.8, 3.6, and 3.4 Å all converged to the same final map resolution

(3.8 Å), and the reconstruction generated with the 3.4 Å cutoff was used for subsequent analysis. Homogeneous reconstruction of the class resembling the reactivation conformation yielded a map from 22,471 particles. The FSC calculated from half-maps reconstructed using HR-HAIR-derived particle poses yielded a nominal resolution of 3.8 Å, although the map visually appears closer to ~4.5-5 Å resolution (Fig. S4A). Following HR-HAIR, both local and NU refinements were performed using a 12 Å initial low-pass filter, with local refinement additionally employing pose/shift Gaussian priors during alignment. Local and NU refinement yielded gold-standard FSC resolutions of 4.6 and 5.1 Å, respectively (Fig. S4B-C). Because the unsharpened HR-HAIR map showed better-defined density, it was selected for restrained model refinement.

HR-HAIR was originally developed to facilitate the reconstruction of very small proteins using only high-resolution information and very small search steps, demonstrating that pose alignment during *ab initio* reconstruction can benefit from incorporating high-resolution information.[7] In our case, keeping the low-resolution information for initial alignment proved beneficial, whereas using only high-resolution information resulted in severe overfitting to noise. Therefore, the only parameters adjusted in the *ab initio* step were the number of classes and maximum resolution cut-off. The synergistic combination of Topaz particle picking, thorough 2D classification, and HR-HAIR enabled the identification and reconstruction of a minor reactivation-state population, while also improving the reconstruction of the dominant resting-state population.

**Cryo-EM Model Building and Refinement**

**Resting-state model:** Structure prediction was performed in Alphafold3[8] using the full-length *T. filiformis* MetH sequence (Uniprot ID: A0A0A2XCD7) with a zinc ion. A starting model was built by individually fitting the Hcy domain, folate domain, and two $B_{12}$ subdomains from the prediction as rigid bodies into the sharpened cryo-EM map in ChimeraX.[9] The modeled zinc was removed due to its unusual predicted position, and both ligands (zinc and cobalamin) were initially placed by aligning to our previous resting-state model (PDB: 8G3H). This initial model was subsequently allowed to relax into the map in ISOLDE[10] (without cobalamin, as ISOLDE does not contain parameterization for it), followed by real space refinement in Phenix[11] after docking back in the cobalamin (global minimization, rigid body, local grid search, anisotropic displacement parameter (ADP), nqh flips morphing and simulated annealing). The model was then migrated back to ISOLDE (again, without cobalamin) for local simulation to help alleviate clashes and fix problematic areas. The cobalamin was then docked back, the position refined by rigid body refinement, and some of the model, especially the areas around the cobalamin, was subsequently manually rebuilt in COOT.[12] The $B_{12}$ restraint file was generated using the Phenix elbow command-line interface with the ideal $B_{12}$ geometry from PDBe-KB ligands. Model statistics, including the model-map FSC, were calculated in Phenix (comprehensive validation, cryo-EM) (Fig. S1, Supplementary Table 1).

**Reactivation-state model:** Alphafold3[8] prediction of the full-length *T. filiformis* MetH's sequence (Uniprot ID: A0A0A2XCD7) consistently yielded a conformation that closely matched the reactivation state, and thus the predicted model was used directly for restrained model refinement against the unsharpened HR-HAIR map. The zinc ion was initially placed by aligning the first two domains to our previous resting-state model (PDB: 8G3H). This initial model was relaxed into its corresponding unsharpened map in ISOLDE[10] using the AlphaFold per-atom pLDDT scores as distance and torsion restraints. These restraints allowed the protein backbone to adjust to the experimental map while preserving reasonable local geometry in regions where the map did not support independent atomic refinement. The cobalamin ligand was then rigid-body docked by aligning the crystal structure of the isolated $B_{12}$-binding domain from *E. coli* MetH (PDB: 1BMT, residues 740-893)[13] to our cryo-EM model (residues 740-872) in ChimeraX.[9] The model was further refined via local grid searching, global minimization, and anisotropic displacement parameter (ADP) refinement over 3 macrocycles in Phenix.[11] To model the cob(II)alamin cofactor, the methyl group was removed from the methylcob(III)alamin coordinates derived from 1BMT, and the ligand was subsequently real-space refined in COOT[12] using restraints based on the ideal geometry of cob(II)alamin. Sidechains were adjusted manually to fix rotamers and clashes. The $B_{12}$ restraint file was generated using the Phenix elbow command-line interface with the ideal $B_{12}$ geometry from PDBe-KB ligands. Model statistics, including the model-map FSC, were calculated in Phenix (comprehensive validation, cryo-EM) (Fig. S4, Supplementary Table 1). To assess potential domain misplacement or over-restraint, we also constructed a conservative comparison model by independently docking individual domains into the local refinement map. This model closely resembled the final model (Cα RMSD of 0.596 Å over all matched pairs), although the latter incorporated ligands and showed higher Q-scores (Fig. S6).

**SAXS of *T. filiformis* MetH**

*T. filiformis* apo-MetH was expressed and purified as previously described[1] and buffer-exchanged into 50 mM HEPES, 150 mM NaCl, 1 mM DTT, pH 7.6 during the final size-exclusion chromatography step. Matching buffer collected from the final size-exclusion chromatography step was used as the SAXS background. SAXS data were collected with an X-ray wavelength of 1.542 Å on a Xenocs BioXolver instrument equipped with a Pilatus3 300K detector. Protein at a concentration of 2 mg/mL was loaded via autoloader into the *in vacuo* sample cell in batch mode at 20 °C, and scattering was collected over a $q$-range of 0.00054-0.44049 $Å^{-1}$, where the momentum transfer variable $q$ is defined as $4\pi/\lambda \sin \theta$, $\lambda$ is the wavelength, and $2\theta$ is the scattering angle. After truncation of the low-$q$ data close to the transmitted beam, the $q$-range used for analyses was 0.0124-0.4405 $Å^{-1}$. Ten individual frames (120 seconds each) were averaged per buffer or protein sample. Data were processed in BioXTAS RAW (Table S2).[14] The pair-distance distribution, $P(r)$, was calculated from the indirect Fourier transform of the scattering intensity $I(q)$ in GNOM.[15] The Cob(II) SAXS data used for comparison in this study was obtained from our previously published work,[1] which was collected and processed using the same experimental procedures.

The apo reactivation model was generated using the refined Cob(II) reactivation-state cryo-EM model. Missing residues in the original model were built using AllosMod-FoXS, and the fit of model to apo protein data was calculated using AllosMod-FoXS with $q$ truncated to 0.25 $Å^{-1}$.[16] Ten models were generated with very similar fitting results, and the one model with the best $\chi^2$ was chosen for further data processing. The fit to the Cob(II) protein data was calculated using the same model as the apo protein data in FoXS[17,18] with $q$ truncated to 0.25 $Å^{-1}$.

**Sequence Selection and Alignment for Four-domain MetH**

All sequences annotated with enzyme commission (EC) number 2.1.1.13 were retrieved from Uniprot[19] (25,549 sequences). Sequences annotated as fragments or lacking proper Uniprot domain annotations were filtered out (20,120 sequences remain). Since cobalamin-dependent activity is of interest, sequences lacking an annotated $B_{12}$-binding domain were removed (16,010 sequences remain). To filter out sequences that contain large insertions or are unlikely to contain more than two intact domains, sequences shorter than 750 residues or longer than 1350 residues were excluded (15,246 sequences remain).

Representative sequences were obtained by grouping redundant sequences with an 85% sequence identity threshold using MMseq2,[20] yielding a non-redundant dataset of 3,704 sequences. Sequences with an annotated Hcy domain shorter than 290 amino acids or an AdoMet domain shorter than 200 amino acids were then excluded, resulting in a final set of 2,205 representative four-domain MetH sequences. Sequence alignment was performed using MAFFT[21] in L-INS-i mode, and sequence logos were generated using WebLogo3.[22]

**Hcy-AdoMet Interface Analysis**

The interfaces between Hcy and AdoMet domains for both the *T. filiformis* Cob(II) MetH reactivation-state model and *T. thermophilus* apo-MetH model (PDB: 8SSC) were analyzed using *qtPisa*.[23–25]

**Supplementary Table 1: Cryo-EM Data Processing and Refinement Statistics**

| | **Resting state** | **Reactivation state** |
|---|---|---|
| EMDB-ID | EMD-78313 | EMD-78314 |
| PDB ID | 37MP | 37MQ |
| **Data processing** | | |
| Microscope | FEI Titan Krios | FEI Titan Krios |
| Camera | Gatan K3 | Gatan K3 |
| Voltage (keV) | 300 | 300 |
| Electron exposure ($e^-/Å^2$) | 61.7 | 61.7 |
| Defocus range (µm) | -0.8 to -2.5 | -0.8 to -2.5 |
| Processing Software | CryoSPARC | CryoSPARC |
| Pixel size (Å) | 1.07 | 1.07 |
| Micrographs collected (used) | 12768 (10341) | 12768 (10341) |
| Final particles (no.) | 113,728 | 22,471 |
| Symmetry imposed | C1 | C1 |
| Map | | |
| Resolution (Å) | 3.4 | 3.8* |
| Resolution range (Å) | 2.8 - 6.6 (75 %) | 3.5 - 9.2 (75%) |
| Sharpening B-factor ($Å^2$) | 113.8 | 38.1 |
| **Model refinement** | | |
| Resolution (Å) (0.143, 0.5) | 3.3, 3.7 | 3.8, 6.4 |
| Model composition | | |
| Non-H atoms | 6530 | 8989 |
| Protein residues | 833 | 1133 |
| Ligands | 2 | 2 |
| B-factors ($Å^2$) | | |
| Protein | 97.96 | 153.26 |
| Ligands | 79.86 | 185.30 |
| R.M.S deviations | | |
| Bond lengths (Å) | 0.012 | 0.006 |
| Bond angles (°) | 1.799 | 1.094 |
| Validation | | |
| *CC* (mask, volume) | 0.70, 0.69 | 0.66, 0.65 |
| Overall Q-score | 0.447 | 0.251 |
| MolProbity score | 1.62 | 1.58 |
| Clashscore | 2.58 | 7.33 |
| Rotamer outliers (%) | 1.80 | 0.44 |
| Ramachandran plot | | |
| Disallowed (%) | 0.00 | 0.00 |
| Allowed (%) | 5.59 | 3.02 |
| Favored (%) | 94.41 | 96.98 |
| Ramachandran Z-scores | | |
| Overall (r.m.s.d) | -2.29 (0.27) | -0.26 (0.24) |
| Helix (r.m.s.d) | -1.86 (0.22) | 0.39 (0.22) |
| Sheet (r.m.s.d) | -1.55 (0.55) | -1.49 (0.46) |
| Loop (r.m.s.d) | -0.83 (0.34) | -0.28 (0.28) |

*Resolution from half-maps reconstructed using HR-HAIR-derived particle poses; the true resolution is ~4.5 - 5 Å. Local refinement yields map (EMD-79288) with a gold-standard FSC resolution of 4.6 Å.

**Supplementary Table 2: SAXS Data Collection Parameters**

| | |
|---|---|
| **Sample details** | |
| Sample | Apo MetH |
| Source organism | *Thermus filiformis* |
| Expression system | *Escherichia coli* |
| *MW* from chemical composition | ~131,143 g $mol^{-1}$ |
| Extinction coefficient (280 nm, theoretical) | 123,080 $M^{-1}$ $cm^{-1}$ |
| Loading concentration | 2 mg/mL |
| Buffer composition | 50 mM HEPES, 150 mM NaCl, 1 mM DTT, pH 7.6 |
| **Data collection parameters** | |
| Instrument/detector | Xenocs BioXolver (Pilatus 300K) |
| Energy (keV) | 8 |
| Detector distance (m) | 0.65 |
| $q$-measurement range (used) ($Å^{-1}$) | 0.00054-0.44049 (0.0124-0.4405) |
| Normalization | Transmitted intensity (direct beam) |
| Exposures | 10 × 120 s |
| Configuration | Batch (oscillating sample) |
| Experimental temperature (°C) | 20 |
| **Data processing/structural parameters** | |
| **Guinier Analysis** | |
| *I(0)* | 0.019 ± 0.00026 |
| $R_g$ (Å) | 38.97 ± 0.78 |
| $qR_g$ range | 0.4832-1.2816 |
| $R^2$ fit | 0.9802 |
| ***P(r)* analysis (GNOM)** | |
| *I(0)* | 0.019 ± 0.00026 |
| $R_g$ (Å) | 40.23 ± 0.96 |
| $D_{max}$ (Å) | 160 |
| $q$-range ($Å^{-1}$) | 0.0124-0.4405 |
| $\chi^2$/total estimate | 1.015/0.797 |
| *MW*, kDa ($V_P$) | 125.5 |

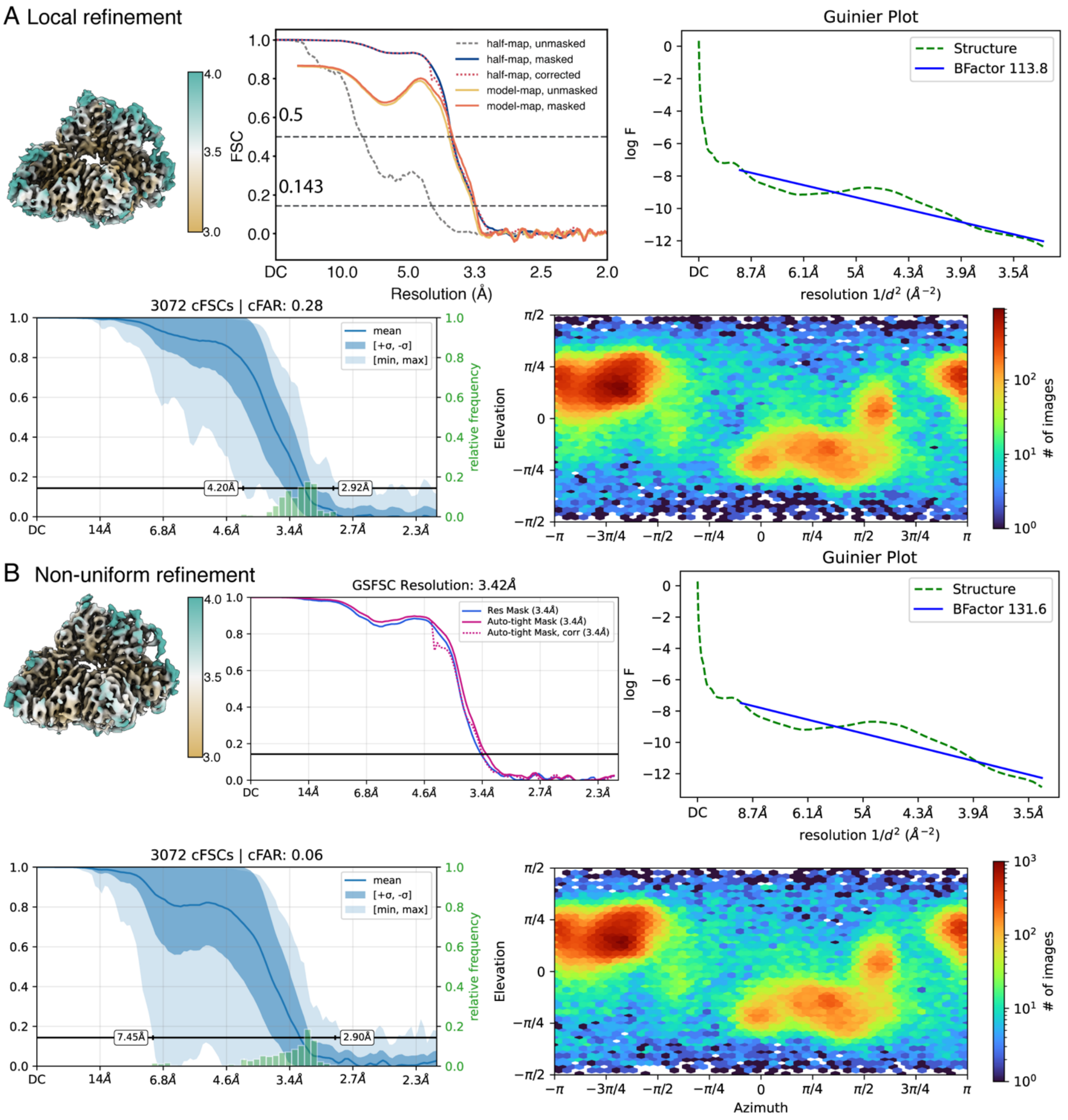


**Figure S1.** Cryo-EM validation of the *T. filiformis* Cob(II) MetH resting-state. Shown are corresponding local resolution (Å) maps, Fourier shell correlation (FSC) curves, B-factor plots, and particle angular distribution for (A) the final map used for model building from local refinement, and (B) the map obtained from non-uniform refinement. Although the two refinements yielded similar estimated resolutions, local refinement produced a more isotropic map, as indicated by the conical Fourier Shell Correlation Area Ratio (cFAR).

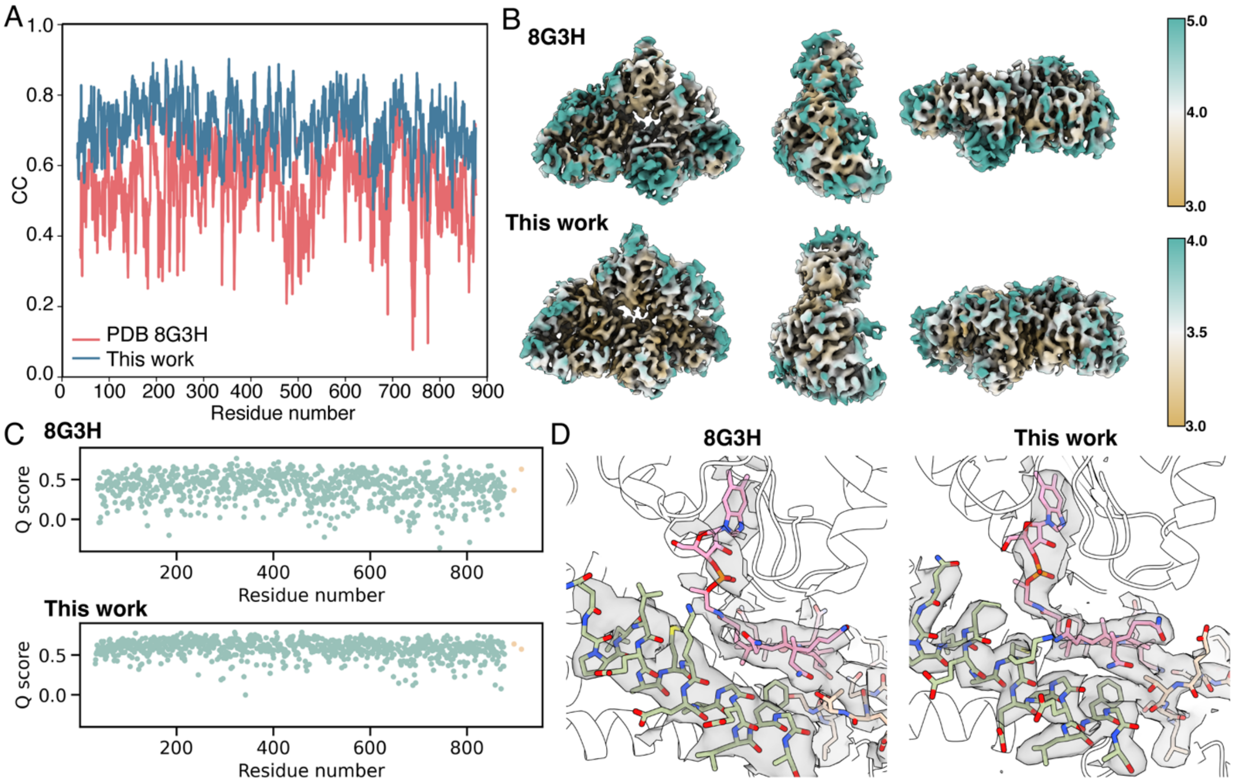


**Figure S2:** Comparison of the previously deposited resting-state structure and map (PDB 8G3H, EMD-29699) with the reprocessed reconstruction and refined model presented in this study. (A) Per-residue cross-correlation for models refined against the original (pink) and reprocessed (blue) maps, showing a ~0.1-0.2 improvement across essentially the entire protein chain for the latter. (B) The new map (bottom) shows uniformly higher local resolution and less anisotropy than the original map (top). (C) Per-residue average Q-score for the original (top) and reprocessed (bottom) maps and corresponding models. Protein residues are colored in green, while cobalamin (left) and zinc (right) are colored in orange. (D) The improved quality of the map can be easily seen in the vicinity of the cobalamin cofactor. In the original map (left), density connecting the corrin ring to the dimethylbenzimidazole tail is not readily visible, while it is clearly resolved in the new map (right). The quality of the new map was also now sufficient to justify refinement of the cofactor itself. The reprocessed map also reveals clear side-chain density for the 697-713 helix in the $B_{12}$-cap subdomain, seen in the foreground (light green) and supports modeling of the 361-370 loop in the folate-binding domain (beige), which was not possible with the original map. The map density is shown carved to within 2.4 Å of the atoms within these regions.

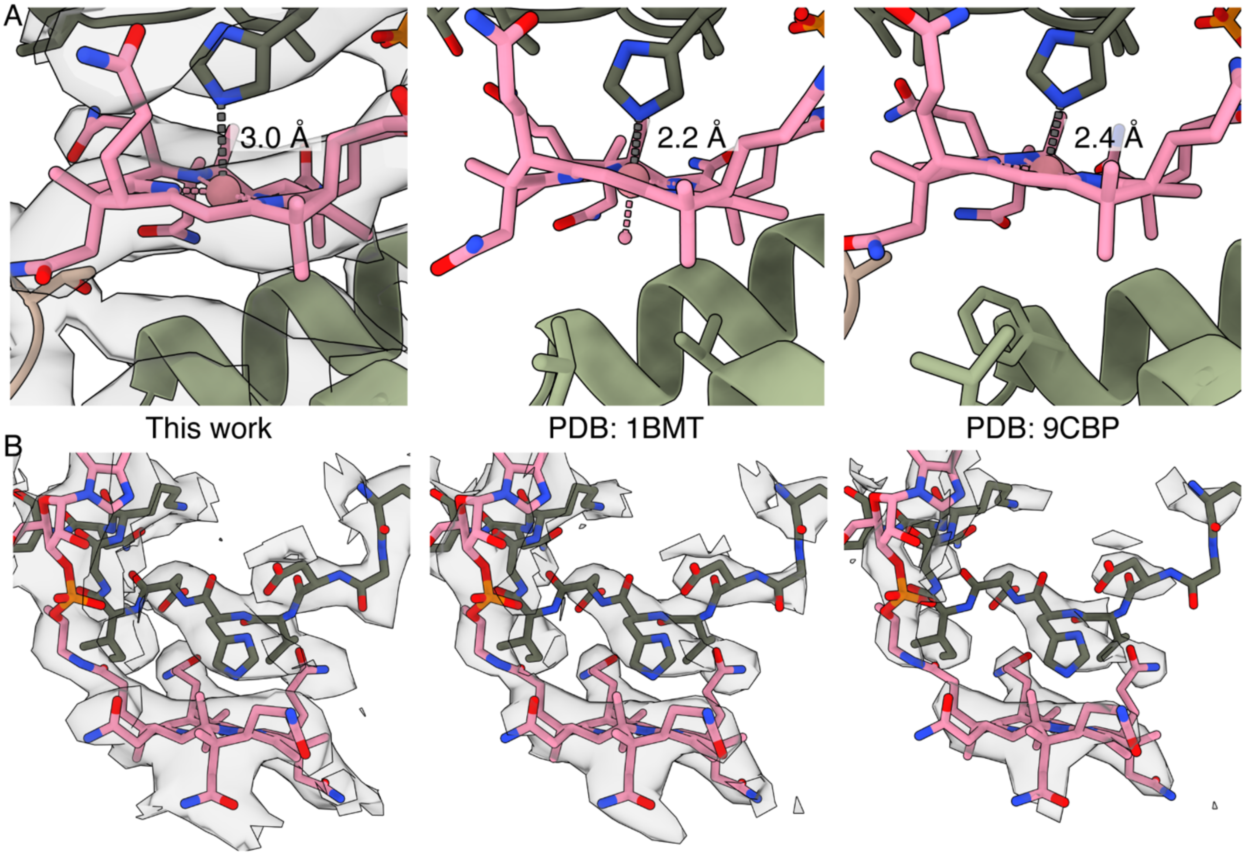


**Figure S3.** Comparison of His-Co bond distances in the cap-on resting-state model obtained in this work with other cap-on MetH structures. (A) The His-Co distance resolved in our refined resting-state model (~3 Å) is longer than what has been observed in previously reported cap-on, His-on crystal structures: ~2.2 Å in PDB: 1BMT, which contained methylcobalamin, and ~2.4 Å in PDB: 9CBP, which contained propylcobalamin. The cryo-EM density in our work is shown at a threshold of 0.213. (B) An alternate view of His759 and cobalamin in our resting-state model with density carved to within 2.5 Å of the shown atoms and displayed at three thresholds: 0.095, 0.15, and 0.213 (left to right).

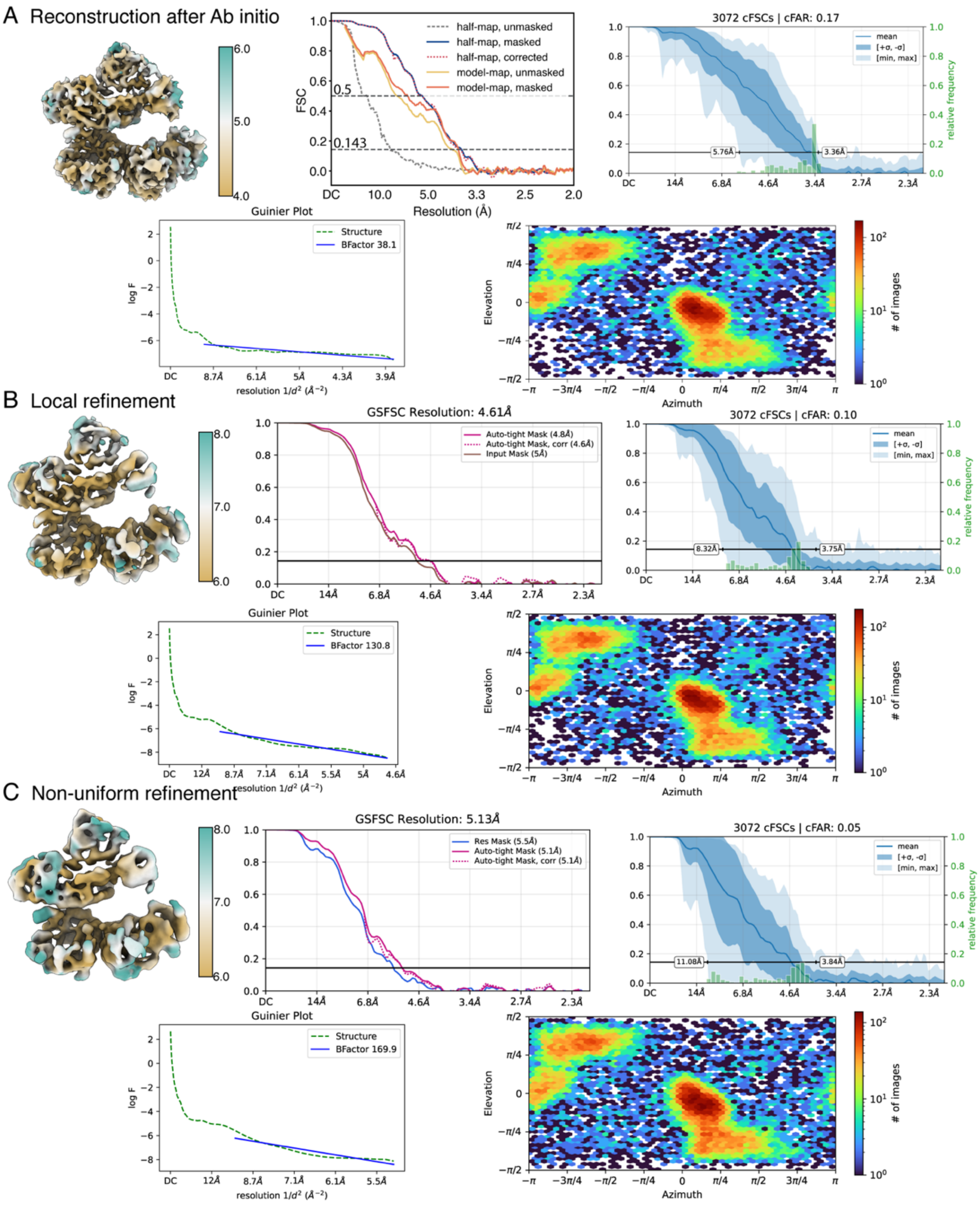


**Figure S4.** Cryo-EM validation of the *T. filiformis* Cob(II) MetH reactivation-state reconstructions. Shown are corresponding local resolution (Å) maps, Fourier shell correlation (FSC) curves, B-factor plots, and particle angular distribution for (A) the final map used for model building, (B) local refinement result and (C) non-uniform refinement result. All three maps show similar overall structural features. Although the HR-HAIR map yields a nominal FSC resolution of 3.8 Å, its apparent resolution is closer to ~4.5–5 Å, consistent with the gold-standard FSC resolutions of 4.6 and 5.1 Å obtained after local and NU refinement, respectively. The HR-HAIR map exhibits better-defined density and was therefore selected for restrained model refinement. The HR-HAIR map is displayed as an unsharpened map, while the remaining two maps are shown sharpened.

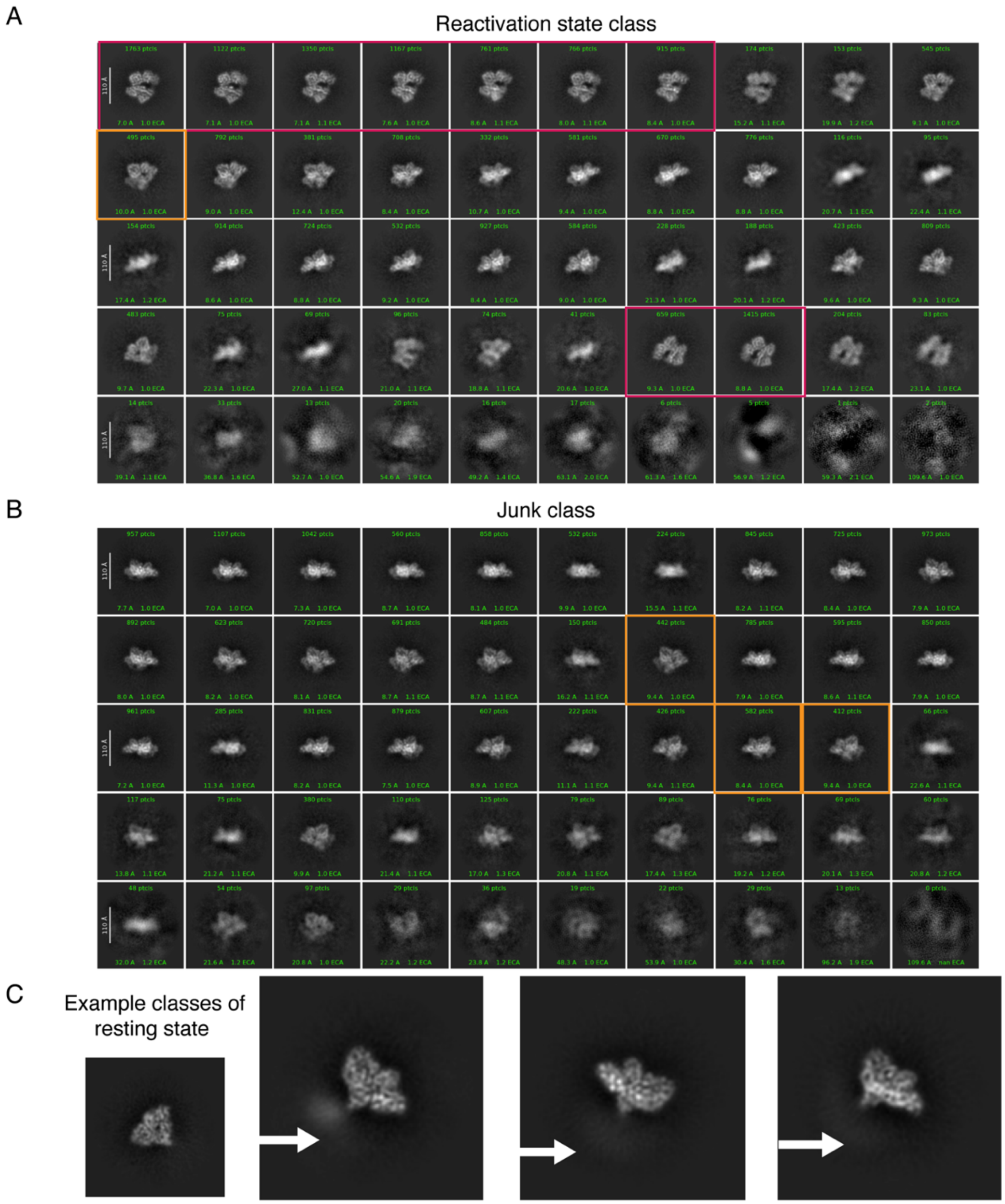


**Figure S5.** Representative 2D classes of particles assigned by HR-HAIR to the reactivation-state and junk classes, compared with representative 2D classes from the resting-state reconstruction. (A) 2D classes of the reactivation-state class contain clear front-view projections of the full-length reactivation conformation (boxed in red), with all four domains visible. (B) 2D classes of the junk particles lack these characteristic views but contain many ambiguous projections of the full-length enzyme. In particular, several resemble the resting state with a stably positioned and resolved AdoMet domain (boxed in orange). (C) In contrast, representative 2D classes from the resting-state reconstruction show well-resolved density for the three N-terminal domains, whereas blurred density (indicated by white arrows) is observed at the nominal position of the AdoMet domain, indicating that it is flexibly linked.

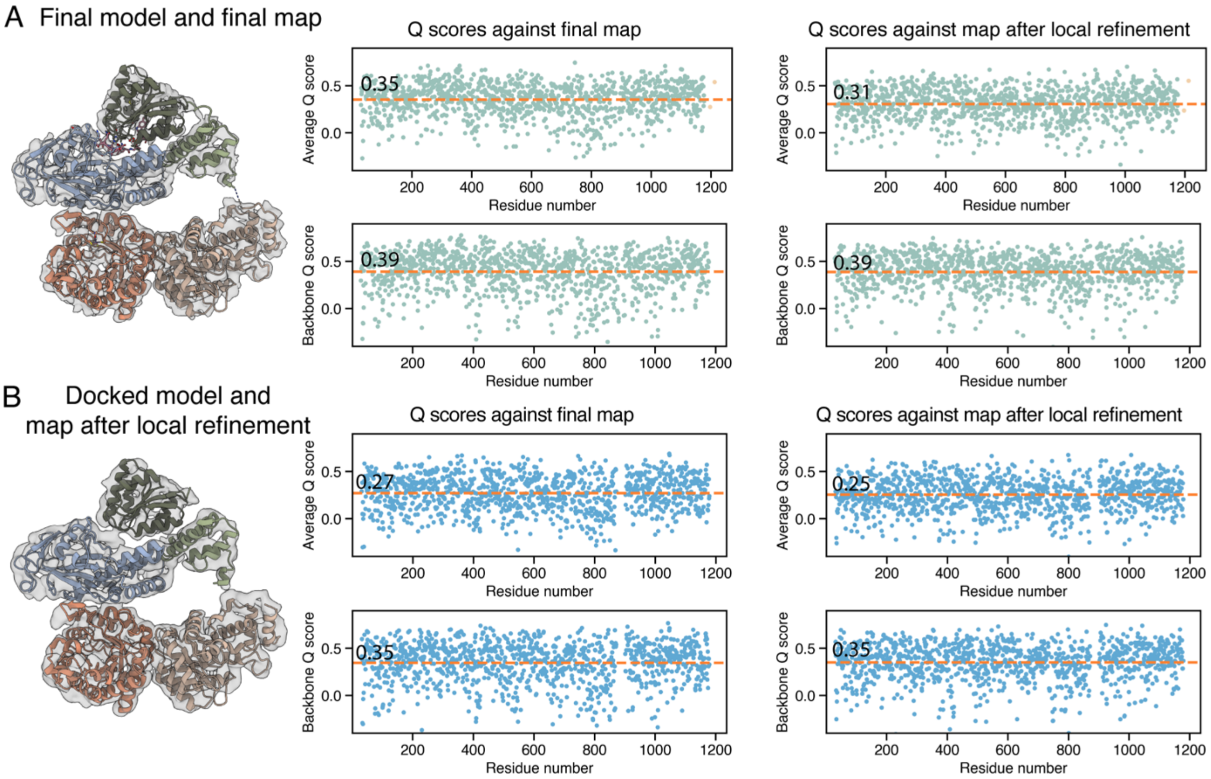


**Figure S6.** Comparison of the final reactivation-state model with a model obtained by per-domain rigid-body docking of an AlphaFold3 prediction. Per-residue average Q-scores and backbone Q-scores calculated in ChimeraX against both the unsharpened HR-HAIR (final) and sharpened local refinement maps are shown for (A) the final model and (B) the per-domain docked model. Protein residues are colored in green or blue, while cobalamin (left) and zinc (right) are colored in orange. The average Q-score across all residues is shown as a dark orange dashed line. The residue ranges used for docking each domain (or sub-domain) are: 31-349, 352-647, 655-738, 744-871, and 901-1178. The final model shows higher Q-scores against both maps, indicating improved agreement with the experimental density. Both models have higher average Q-scores against the HR-HAIR map than against the local refinement map.

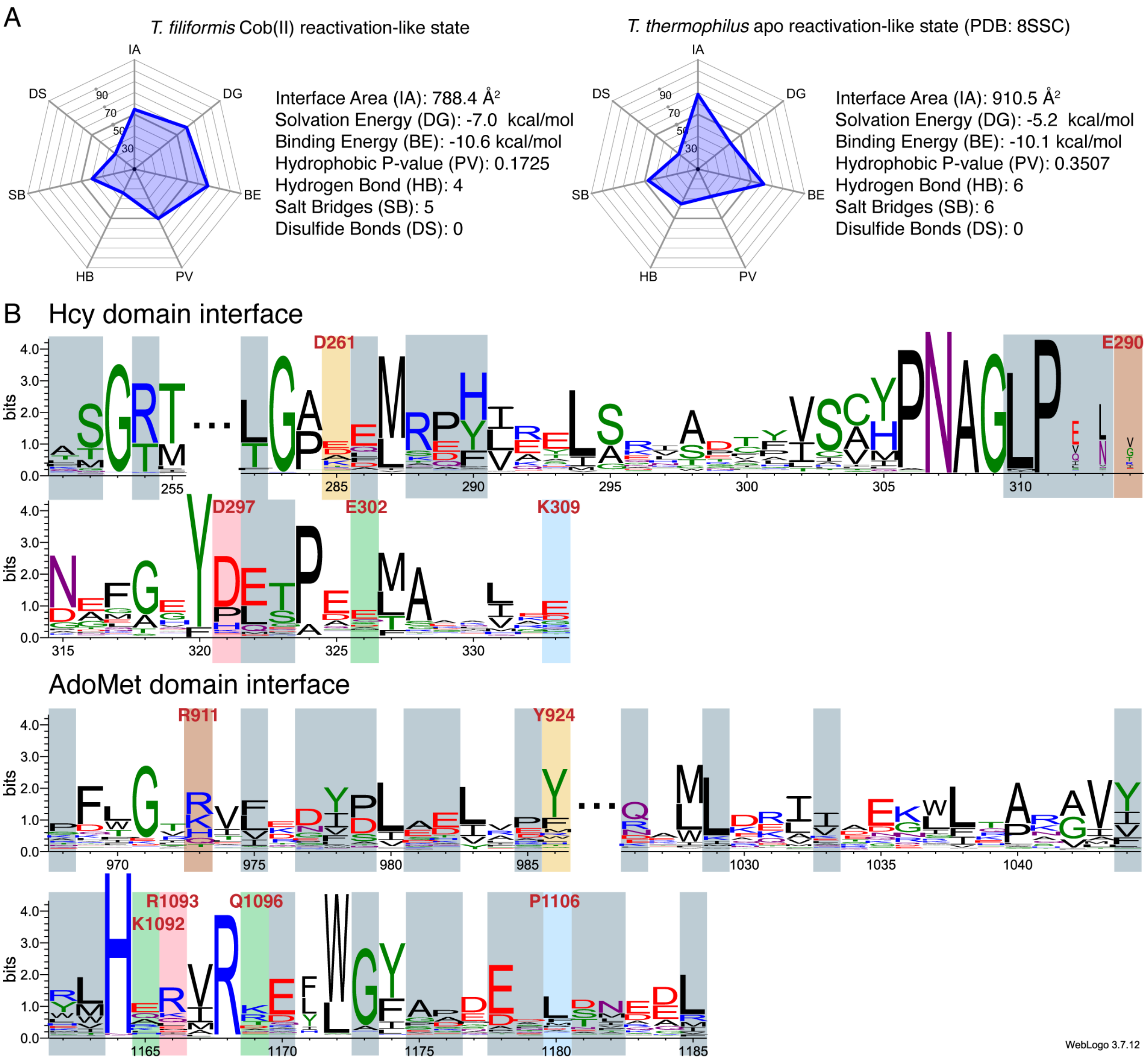


**Figure S7.** Hcy-AdoMet domain interface analysis. (A) *QtPisa* shows that the Hcy-AdoMet interfaces in both the *T. filiformis* Cob(II) reactivation conformation (this work) and the *T. thermophilus* apo-MetH conformation (PDB: 8SSC) are similar and moderately strong. (B) Sequence logos of 2,205 representative MetH sequences show moderate conservation for interfacial residues that participate in non-electrostatic interactions (gray). Residues involved in electrostatic interactions (colored pairs) are less well-conserved.

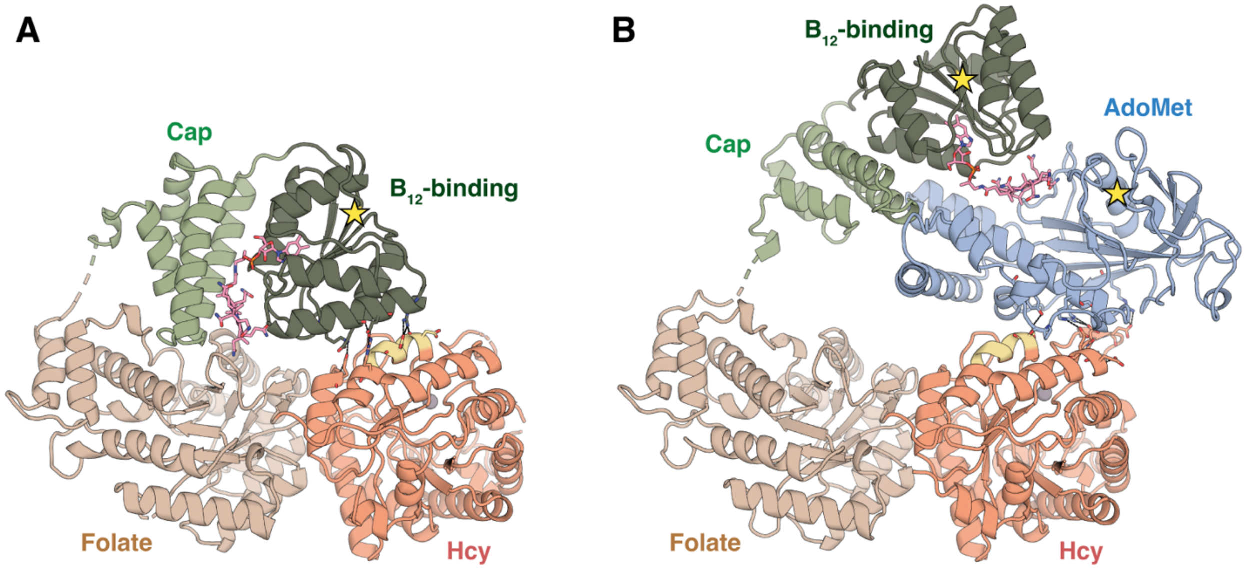


**Figure S8.** The resting and reactivation conformations of *T. filiformis* MetH are distinct states exhibiting mutually exclusive interactions with the Hcy domain. (A) The resting-state model and (B) the reactivation-state model share a partially overlapping interaction surface on the Hcy domain (yellow). Comparison of the two models, aligned on the Hcy domain, reveals that the $B_{12}$-binding subdomain and AdoMet domain occupy mutually exclusive positions relative to the Hcy domain, although the specific polar contacts are not conserved. Interconversion between these states requires substantial structural rearrangements, including disengagement of the $B_{12}$-binding subdomain from the Hcy domain, accommodation of the AdoMet domain at this interface, and uncapping of the $B_{12}$ domain to interact with the AdoMet domain. Between the two conformations, the center of mass of the $B_{12}$-binding subdomain is displaced by more than 30 Å, while the center of the cobalamin corrin ring moves approximately 40 Å. In the resting state, the AdoMet domain is highly mobile and is not resolved in a single position; the yellow star in panel A marks the last modeled residue of the $B_{12}$ domain and the point from which the flexible linker to the AdoMet domain extends. In the reactivation state, the AdoMet domain adopts a well-defined position, while the connecting linker remains flexible; the yellow stars in panel B mark the modeled residues flanking this unresolved linker and illustrate the connectivity that permits the large domain rearrangements between the two states.
